# Genetic requirements of *Mycobacterium tuberculosis* for survival under pathogen-specific immunity

**DOI:** 10.1101/2024.09.26.615000

**Authors:** Kimra S. James, Neharika Jain, Kelly Witzl, Nico Cicchetti, Sarah M. Fortune, Thomas R. Ioerger, Amanda J. Martinot, Allison F. Carey

**Affiliations:** Division of Microbiology & Immunology, Department of Pathology, University of Utah, Salt Lake City, UT; Department of Infectious Disease and Global Health, Tufts University Cummings School of Veterinary Medicine, North Grafton, MA; Department of Immunology & Infectious Disease, Harvard T.H. Chan School of Public Health, Boston, MA; Department of Computer Science, Texas A&M University, College Station, TX

## Abstract

*Mycobacterium tuberculosis* (Mtb), the etiologic agent of tuberculosis (TB), remains a persistent global health challenge due to the lack of an effective vaccine. The only licensed TB vaccine, Bacille Calmette-Guerin (BCG), is a live attenuated strain of *Mycobacterium bovis*, that protects young children from severe disease but fails to provide protection through adulthood. It is unclear why BCG-induced immunity provides incomplete protection despite inducing a robust CD4+ T cell response, a known critical correlate of protection against Mtb. We set out to interrogate mycobacterial determinants of vaccine escape using a functional genomics approach, TnSeq, to define mycobacterial genes required for survival in mice vaccinated with BCG, the live attenuated Mtb vaccine, ΔLprG, and in mice with Mtb immunity conferred by prior infection. We find that critical virulence genes associated with acute infection and exponential growth are less essential in hosts with adaptive immunity, including genes encoding the Esx-1 and Mce1 systems. Genetic requirements for Mtb growth in vaccinated and previously Mtb-infected hosts mirror the genetic requirements reported for bacteria grown under *in vitro* conditions that reflect aspects of the adaptive immune response as well as in mouse genetic backgrounds where bacterial growth is restricted. Across distinct immunization conditions, differences in genetic requirements between live attenuated vaccines and vaccination routes are observed, suggesting that different immunization strategies impose distinct bacterial stressors. Collectively, these data support the idea that Mtb requires genes that enable stress adaptation and growth arrest upon encountering the restrictive host environment induced by the adaptive immune response. We demonstrate that TnSeq can be used to understand the bacterial genetic requirements for survival in vaccinated hosts across pre-clinical live attenuated vaccines and therefore may be applied to other vaccine modalities. Understanding how Mtb survives in the setting of vaccine-induced immunity could uncover bacterial vulnerabilities to inform the development and prioritization of new vaccines or adjuvant therapies.

## INTRODUCTION

*Mycobacterium tuberculosis* (Mtb), the etiologic agent of tuberculosis (TB), caused an estimated 10 million infections in 2021, resulting in over 1.5 million deaths across the globe^1^. One reason why TB remains a persistent global health challenge is the lack of an effective vaccine. The only licensed TB vaccine, Bacille Calmette-Guerin (BCG), is a live attenuated strain of *Mycobacterium bovis* developed over 100 years ago. BCG is effective in preventing severe, extrapulmonary TB disease in infants and young children, a major cause of morbidity and mortality in TB endemic regions. However, BCG provides minimal protection against pulmonary TB in adults, the most common form of the disease^2^.

It is unclear why BCG-induced immunity provides incomplete protection. BCG elicits a robust CD4^+^ T-cell response, which, via IFNγ production, enhances macrophage oxidative burst and phagolysosomal maturation, resulting in increased bacterial killing^3^. CD4^+^ T-cells also promote antibody production and maturation^4^, and recruitment and activation of cytotoxic CD8^+^ T cells and other effector cells^5^. While IFNγ production is clearly necessary for vaccine-induced protection, the frequency of IFNγ producing T-cells in vaccinated individuals is not predictive of disease risk, suggesting that additional bacterial control mechanisms are involved^6^. In contrast to the intensive studies of the immune response induced by BCG vaccination, mycobacterial determinants of vaccine escape remain unknown, further hampering the rational design of improved immunization strategies. Understanding how Mtb survives in the setting of vaccine-induced immunity could uncover bacterial vulnerabilities to inform the development and prioritization of new vaccines or adjuvant therapies.

With the goal of uncovering such vulnerabilities, we used a functional genomics approach, TnSeq, to define mycobacterial genes required for survival in vaccinated mice and mice with immunity to prior Mtb infection. TnSeq entails selecting a saturated transposon insertion library under a defined growth condition, followed by transposon junction sequencing for insertion mapping. The distribution of insertions across the genome is compared between output and control libraries to identify genes with differences in transposon insertion frequency. Genes with fewer insertions in the output library are considered conditionally essential for bacterial growth, while genes with more insertions are considered conditionally non-essential. TnSeq has been used to define mycobacterial genes required to survive under numerous *in vitro* and *in vivo* conditions, providing important insights into bacterial physiology^7–16^. We therefore reasoned that TnSeq could be used to understand how Mtb survives the numerous pressures imposed by the pathogen-specific adaptive immune response.

We find that critical virulence genes associated with acute infection and exponential growth are less essential in hosts with adaptive immunity, including genes encoding the Esx-1 and Mce1 systems. This suggests that the cellular microenvironment of the vaccine-induced immune response is more akin to what the bacteria experience during chronic infection, when bacterial growth becomes constrained. Consistent with this model, genetic requirements for Mtb growth in vaccinated and previously Mtb-infected hosts mirror the genetic requirements reported for bacteria grown under *in vitro* conditions that reflect aspects of the adaptive immune response, for example, the low pH of the mature phagolysosome, as well as in mouse genetic backgrounds where bacterial growth is restricted. Across distinct immunization conditions, differences in genetic requirements between live attenuated vaccines and vaccination routes are observed, suggesting that different immunization strategies impose distinct stressors.

Collectively, these data support the idea that Mtb requires genes that enable stress adaptation and growth arrest upon encountering the restrictive host environment induced by the adaptive immune response. Understanding the bacterial genetic requirements for survival in vaccinated hosts across pre-clinical live attenuated vaccines may lead to a more nuanced understanding of correlates and mechanisms of protection and lead to improved vaccines to curb this ongoing global health emergency.

## RESULTS

### Comprehensively defining *M. tuberculosis* genetic requirements for survival in BCG vaccinated hosts

The added immune pressure conferred by BCG vaccination is expected to impose substantial selection against the bacterium during infection, yet the genes and pathways Mtb relies on to survive this selective pressure remain largely unknown. To define this suite of bacterial genes, we infected BCG-vaccinated and age-matched naïve C57BL/6 mice with a saturated transposon library generated in the widely-used Mtb reference strain H37Rv (Figure 1A). Two- and four-weeks post-challenge, mice were sacrificed and tissue harvested for CFU enumeration and to recover genomic DNA from the surviving mutants for transposon-junction sequencing. Subcutaneous BCG vaccination reduced bacterial burden by approximately 1.5 log at these time points, consistent with prior studies on the efficacy of BCG in the mouse model of Mtb infection (7.054 and 5.511 log_10_ CFU/spleen in naïve and BCG-vaccinated mice, respectively, two weeks post-challenge; 6.958 and 5.264 log_10_ CFU/spleen in naïve and BCG-vaccinated mice four weeks post-challenge). To identify genes that are differentially essential between naïve and BCG vaccinated animals, we performed head-to-head comparisons of the output transposon libraries using the Transit resampling method^8^. With correction for multiple testing, we found that 102 genes were differentially essential two weeks post-challenge, and 64 genes were differentially essential four weeks post-challenge (adj. p-value <0.05, Figure 1B, Supplemental Table 1).

**Figure 1.**
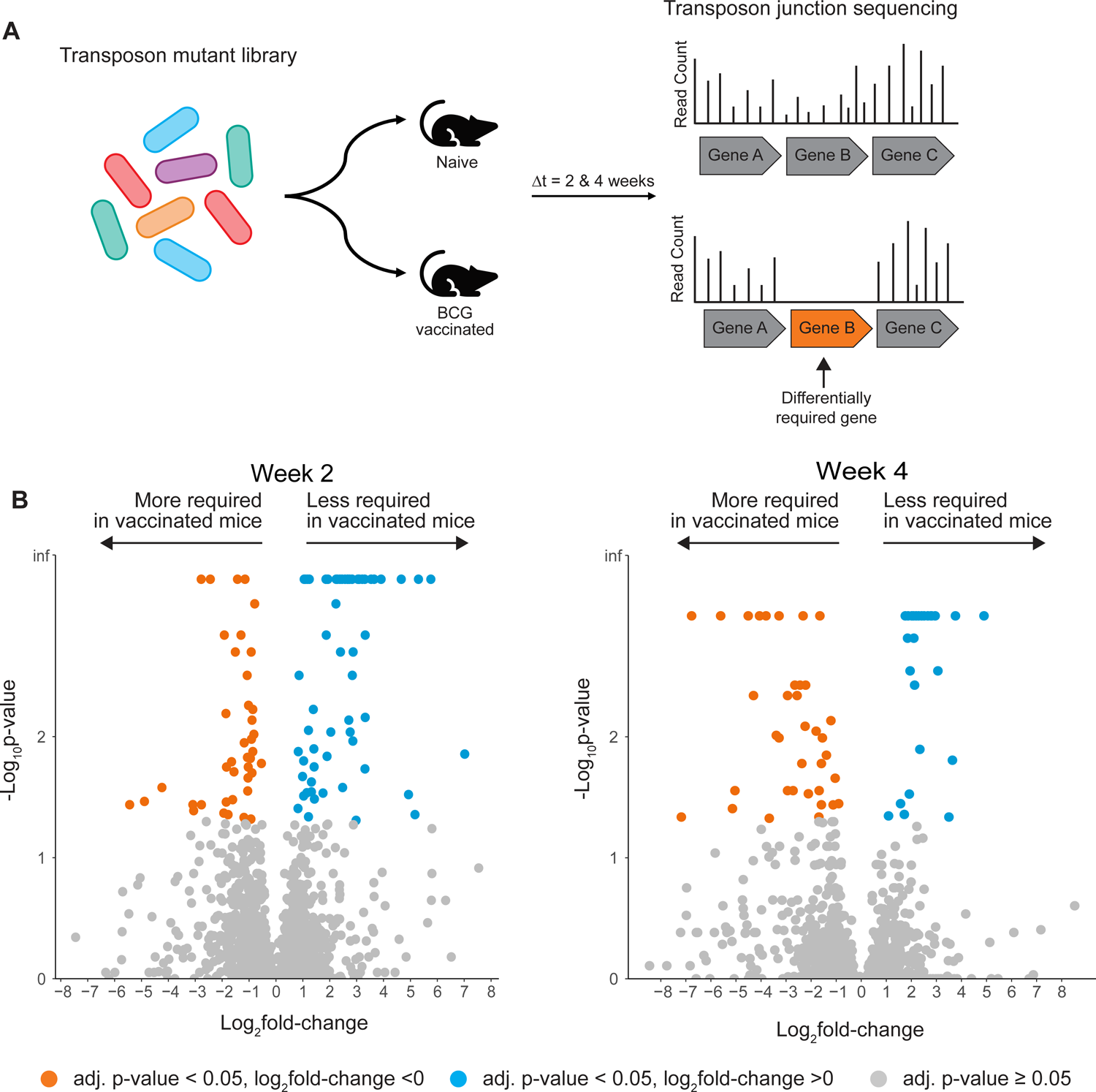
Mtb genes required for survival and growth in BCG vaccinated mice. A) Schematic of experimental strategy for TnSeq screen. B) Volcano plots highlighting differentially required genes in BCG vaccinated mice relative to naïve mice two weeks (left panel) and four weeks (right panel) post-infection.

Given the selective pressure imposed by vaccination, we expected to identify genes that were more essential in immunized relative to naïve mice. Consistent with this expectation, 41 genes had a statistically significant decrease in transposon read count in BCG vaccinated mice compared to naïve mice two weeks post-challenge. The set of genes conditionally essential for survival in vaccinated mice are distributed across a number of functional categories, and pathway analysis did not identify significant enrichment in any category (Supplemental Figure 1A). However, among the most differentially essential genes are *katG/Rv1908c*, which encodes a catalase-peroxidase that detoxifies reactive oxygen species; *mosR/Rv0348*, a transcriptional regulator induced by hypoxia and one of the most highly expressed genes during chronic infection in mice; *virS/Rv3082c*, a pH-responsive transcriptional regulator activated in the phagolysosome; and *ctpC/Rv3270*, a P-type ATPase responsible for metalating *sodA*, the major superoxide dismutase in Mtb. Altogether, these genes represent responses to host-imposed stresses enhanced by BCG vaccination, such as the CD4^+^ T-cell mediated increase in IFN-γ and TNF-α from infected and bystander macrophages^10^.

Four weeks post-challenge, 36 genes were significantly more required in vaccinated as compared to naïve mice. Only one gene, the catalase-peroxidase *katG/Rv1908c*, overlapped with the more essential genes at the two week time point (Supplemental Figure 1A), suggesting continued, enhanced oxidative stress in vaccinated hosts. Among the other genes most essential for growth four weeks post-challenge are *rsfB*/*Rv3687c*, an anti-anti-sigma factor that enhances the activity of the alternative sigma factor 0^F^, which plays a central role in stress response and persistence during chronic infection^17^; *cuvA/Rv1422*, which encodes a protein involved in lipid utilization and has a persistence defect when deleted^18^; *fadD6/Rv1206*, an acyl-CoA synthetase that promotes triacylglycerol accumulation and mycobacterial dormancy^19^; and *Rv2628*, a hypothetical protein in the DosR dormancy regulon that encodes an antigen associated with latent infection^20,21^. Altogether, many of these genes are linked to induction and maintenance of Mtb dormancy, suggesting that Mtb’s ability to arrest growth is critical to surviving the memory immune response.

### Canonical virulence genes are less required in BCG vaccinated mice

Unexpectedly, approximately half of the statistically significant genes were less essential in BCG vaccinated mice compared to naïve mice, displaying an increase in transposon insertion count: 61 genes two weeks post-challenge and 28 genes four weeks post-challenge (Figure 1B, Supplemental Table 1). These two gene sets are highly overlapping, and pathway analysis found that the functional category ‘virulence’ was significantly over-represented at both time points (Supplemental Figure 1B). Notably, many of the less essential genes belong to two systems that play key roles in Mtb pathogenesis and its intracellular lifestyle: the Esx-1 Type VII secretion system, and the Mce1 lipid uptake system (Figure 2A, Supplemental Figure 1B). Both the Esx-1 and Mce1 systems are known to be essential for Mtb growth *in vivo*^15,22–25^.

**Figure 2.**
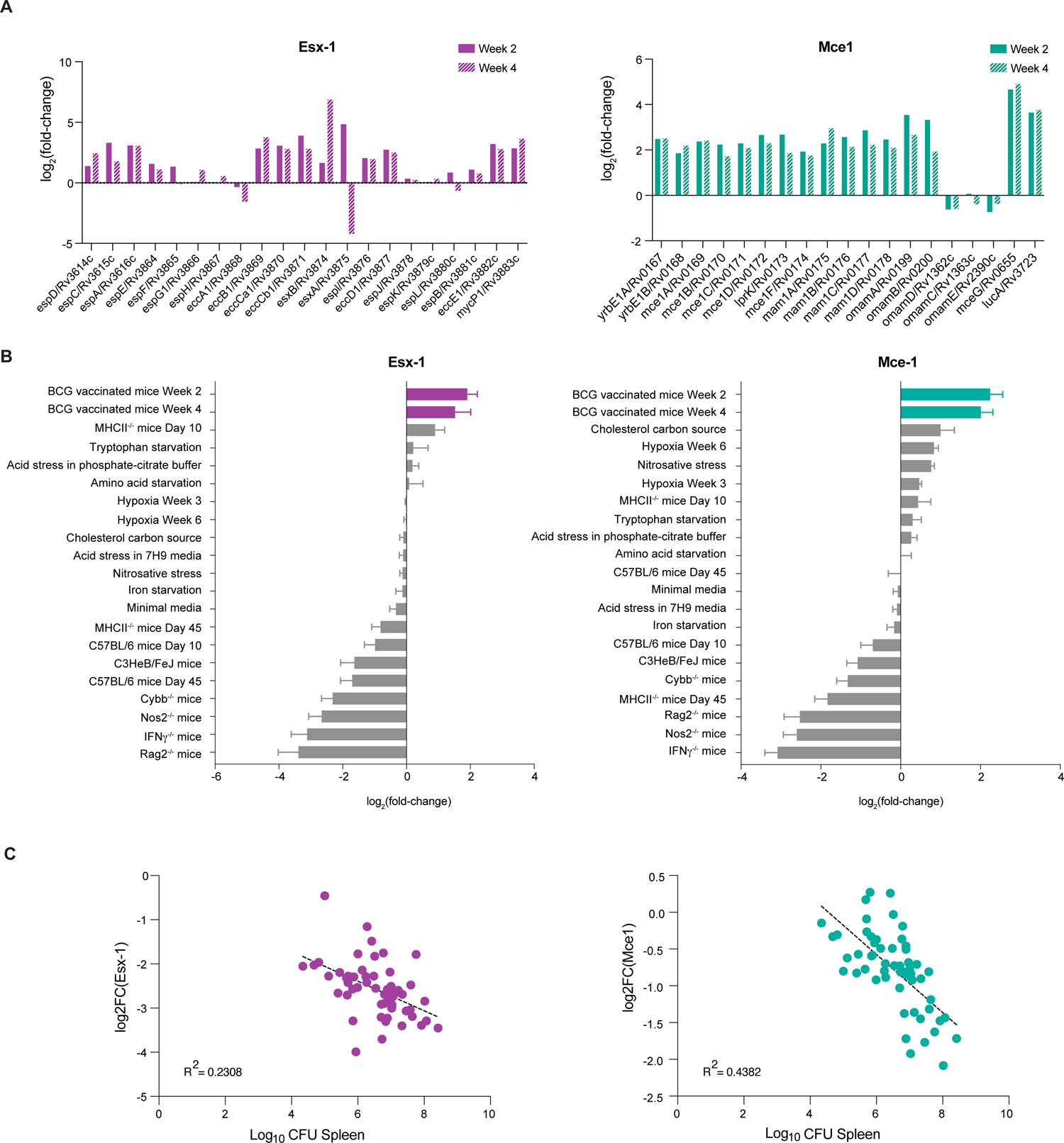
Critical virulence genes are less essential in BCG vaccinated mice. A) Log_2_(fold-change) in transposon read count for each gene in the Esx-1 system and Mce1 system in BCG vaccinated mice relative to naïve mice. B) Mean log_2_(fold-change) in transposon read count for genes in the Esx-1 or Mce1 systems across defined *in vitro* conditions and mouse genetic backgrounds. Error bars represent SEM. Data obtained from MtbTnDb database. C) Correlation between log_2_(fold-change) in transposon read count for Esx-1 or Mce1 genes and bacterial burden across a set of genetically diverse mouse backgrounds.

The Esx-1 system consists of a Type VII transmembrane secretion system and secreted effector proteins which are critical to Mtb pathogenesis. A key role of Esx-1 is to permeabilize the phagolysosomal membrane, facilitating release of bacterial components into the cytosol, thereby initiating a complex innate immune response from the host^26–31^. The Esx-1 secretion system, secreted proteins, and regulatory factors are encoded across two unlinked genomic loci^32^. Genes across both loci were significantly less essential in BCG vaccinated mice as compared to naïve mice at both time points (Figure 2A, Supplemental Figure 2A, Supplemental Table 1). Most genes in the Esx-1 loci that did not reach statistical significance had a non-significant increase in the number of transposon insertions (Figure 2A, Supplemental Figure 2A, Supplemental Table 1).

Mce1 is also a transmembrane complex, responsible for importing host-derived fatty acids, an important nutrient for Mtb during acute infection^23^. The Mce1 complex is encoded across a 12-gene operon, with seven accessory genes located elsewhere across the genome. All 12 of the genes in the Mce1 operon, in addition to the accessory genes *lucA, omamA*, and *mceG,* were significantly less essential in BCG vaccinated mice compared to naïve mice at one or both time points (Figure 2A, Supplemental Figure 2A, Supplemental Table 1).

We note that the reduced essentiality of Esx-1 in BCG vaccinated mice is relative, rather than absolute. Transposon insertions across the Esx-1 genes are over-represented in vaccinated as compared to naïve mice. However, relative to the input library, they are under-represented (Supplemental Figure 2B). By contrast, bacteria with transposon insertions across the Mce1 genes are more abundant in vaccinated mice as compared to the input library (Supplemental Figure 2B), suggesting an absolute growth advantage for Mce1 mutants in vaccinated mice.

To validate our observation that the Esx-1 virulence system is less essential in vaccinated mice, we challenged groups of BCG vaccinated and naïve animals with either wild-type H37Rv or an Esx-1 deletion mutant^28^ (Supplemental Figure 2C). Two- and four-weeks post-challenge, animals were sacrificed and CFU enumerated from the spleens and lungs. To quantify the relative fitness cost of Esx-1 disruption in naïve compared to BCG-vaccinated animals, we calculated the log_10_ difference in bacterial burden (CFU) between naïve and vaccinated animals. We found that the Esx-1 deletion strain had a substantial growth defect in naïve mice two and four weeks post-infection, consistent with previous reports^15,26^. However, this growth defect was reduced in BCG-vaccinated mice in the spleen, with an even greater effect in the lung (Supplemental Figure 2C). These results support our TnSeq findings that the *in vivo* essentiality of a canonical virulence factor, Esx-1, is partially rescued in the context of BCG vaccination.

### Decreased essentiality of Esx-1 and Mce1 genes corresponds to conditions promoting Mtb growth arrest

What does it mean for virulence systems to be less essential in vaccinated mice? To gain biological insight, we evaluated the requirement for Esx-1 and Mce1 genes across TnSeq datasets from the MtbTnDb database^33^. MtbTnDb includes data from TnSeq experiments across numerous growth conditions, and the raw read data have been standardized and analyzed by Transit. We extracted data from MtbTnDb from wild-type H37Rv transposon libraries selected under defined *in vitro* conditions or in defined knockout mice for comparison (Supplemental Table 2). We then plotted the mean log_2_fold-change in transposon read count averaged across Esx-1 or Mce1 genes for each dataset to identify conditions that phenocopy the bacterial requirement for these genes during vaccination (Figure 2B). As expected, Esx-1 genes are essential in wild-type, C57BL/6 mice as compared to the *in vitro* selected input library at days 10 and 45 post-infection (Figure 2B). Esx-1 genes are also conditionally essential in immune-deficient mice, including Rag, NOS, IFNγ, and Phox (Cybb) knockout backgrounds. By contrast, conditions where Esx-1 is not essential include *in vitro* hypoxia, *in vitro* acid stress, and *in vitro* amino acid limitation.

Evaluating the essentiality of Mce1 genes across these same conditions finds that Mce1 genes are not required when grown *in vitro* in cholesterol media as compared to glycerol media, consistent with the known role of Mce1 as a fatty-acid importer^34^. This suggests that bacteria surviving the immune pressure of vaccination may have distinct metabolic activity or access to different nutrient sources. Overall, the rank order of conditions for Mce1 genes is similar to that for Esx-1 genes. Like Esx-1 genes, Mce1 genes are less essential under *in vitro* hypoxia, *in vitro* acid stress, and *in vitro* amino acid limitation, and more essential in Rag, NOS, IFNγ, and Phox knockout mice. Because bacterial replication is uncontrolled in these immune-deficient mouse backgrounds, we hypothesized that the requirement for Esx-1 and Mce1 may reflect rapid bacterial growth.

To test this model, we analyzed a large, published dataset of bacterial burden and TnSeq measurements across the collaborative cross mice, a panel of genetically diverse mice with variable capacity for bacterial control^10,35^. We found bacterial burden (CFU) to be negatively correlated with the log_2_fold-change in transposon count across Esx-1 genes (R^2^ = 0.2308) and the Mce1 genes (R^2^ = 0.4382) (Figure 2C, Supplemental Table 3). This is consistent with a model in which Esx-1 and Mce1 are required under conditions of rapid bacterial replication, and the decreased essentiality of these gene sets in vaccinated mice may reflect a non-replicating state of the surviving bacteria.

### Differential effect of immunization strategy on essentiality of Esx-1 and Mce1

Given the poor performance of BCG in conferring lasting, protective immunity against Mtb infection and disease, significant efforts have been devoted towards the development of improved vaccine strategies. These include subunit vaccines, novel attenuated whole cell vaccines, and methods of delivery that promise to enhance the immune response, including mucosal and intravenous routes of administration. Quantitative and qualitative differences in immune response and degree and duration of protection have been described under various vaccination strategies, yet bacterial physiology under these conditions has not been interrogated.

We therefore expanded our vaccine TnSeq challenge model to include additional immunization conditions (Figure 3A). We included mice vaccinated with BCG via intravenous route (BCG-IV), which has been shown to confer increased protection against Mtb compared to BCG-SQ in non-human primates and to enhance trained immunity in murine models. In non-human primates, BCG-IV induces greater numbers of antigen-specific T-cells, and in mice, BCG-IV reprograms the myeloid hematopoietic compartment ^36–38^. We also included mice vaccinated with a live, attenuated strain of Mtb, ΔLrpG, which has been shown to confer greater protection than subcutaneous BCG (BCG-SQ) against Mtb in the C3HeB/FeJ mouse model of infection, induce a more robust cytokine response, and elicit a polyfunctional CD4^+^ T-cell response^39,40^. Finally, we included mice that had been infected via aerosol route with H37Rv then cured with combination antibiotic therapy (Mtb+cure). This approximates reinfection in humans and may provide insight into how Mtb escapes the immunity conferred by natural infection. A meta-analysis of epidemiological studies from the pre-antibiotic era found that prior Mtb infection confers a degree of protection against TB disease similar to that of BCG vaccination^41^, while ongoing Mtb infection confers near-sterilizing protection against secondary infection in a non-human primate model^42^. In a mouse model, prior Mtb infection and cure afforded a log reduction in Mtb burden in mice, similar to the level of protection afforded by subcutaneous BCG vaccination, the traditional benchmark for TB vaccine evaluation in pre-clinical mouse models^43^.

**Figure 3.**
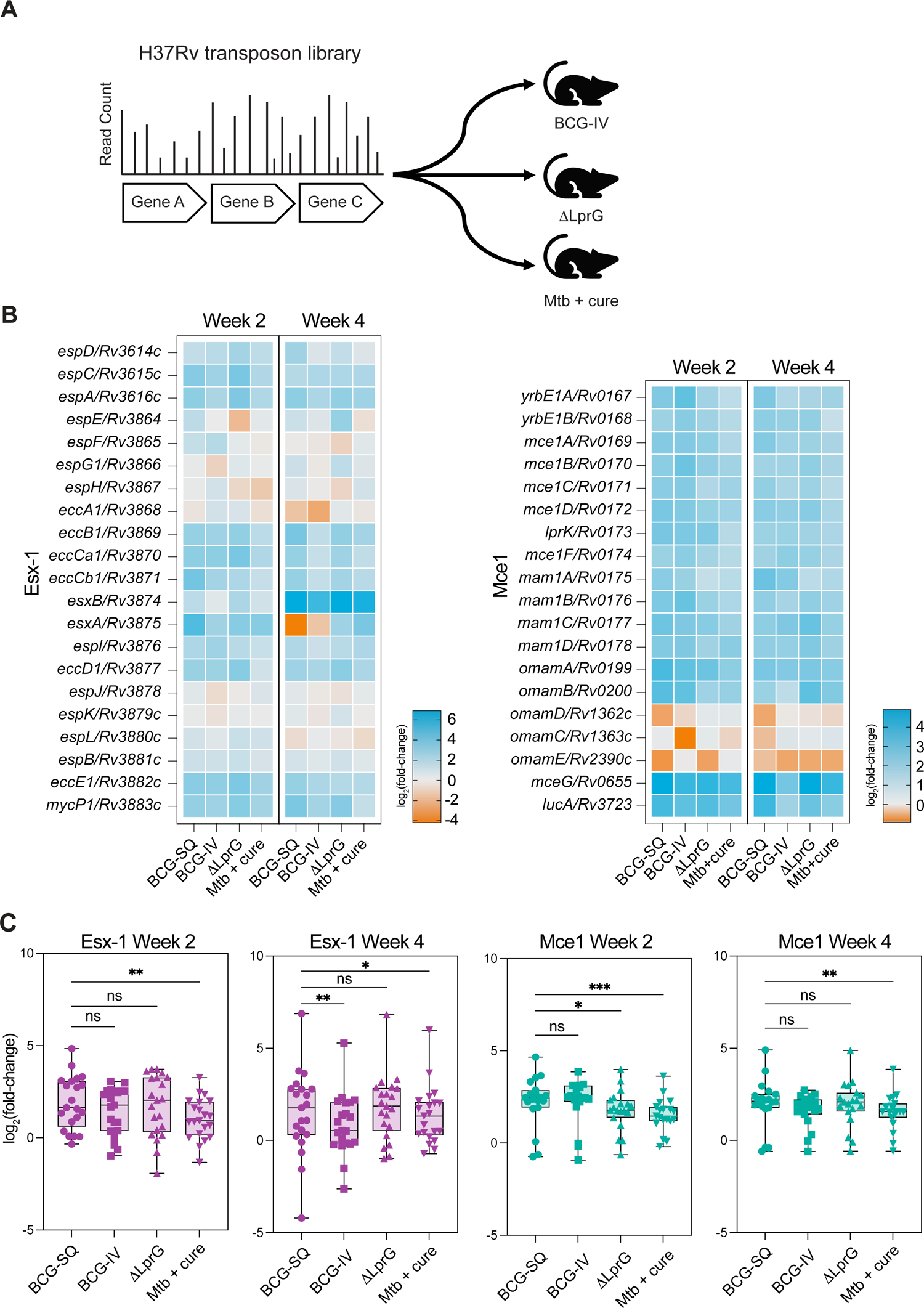
Genetic requirements across distinct immunization conditions. A) Experimental strategy for the expanded TnSeq immunization screen. B) Heatmaps of log_2_(fold-change) in transposon read counts across Esx-1 and Mce1 genes across immunization conditions relative to naïve mice. C) Box plots showing log_2_(fold-change) in transposon read counts across Esx-1 and Mce1 genes. Significance determined by Friedman test with Dunn’s multiple comparison to BCG-SQ condition.

Mice in each group were challenged with the H37Rv transposon library and surviving mutants recovered and sequenced as described above. We found that bacterial burden was reduced by approximately 1.5 logs in each arm of the experiment compared to naïve animals, and was not significantly different across arms (Supplemental Figure 3A). While this suggests quantitatively similar selective pressure across the immunization strategies, we wanted to assess whether there were qualitative differences among these conditions, as reflected in the bacterial genes essential for growth. Given the striking difference in essentiality of Esx-1 and Mce1 genes in BCG-SQ vaccinated as compared to naïve animals, we first evaluated the essentiality of these systems.

Compared to naïve mice, many genes in the Esx-1 and Mce1 loci were significantly less essential in the ΔLrpG, BCG-IV, and Mtb+cure arms, consistent with our observations in BCG-SQ vaccinated animals (Supplemental Table 1). However, there are quantitative differences in the essentiality of these genes across the vaccination strategies (Figure 3B). When transposon read counts are averaged across all genes in the Esx-1 system, there is a significant difference among the vaccine strategies at both time points (p = 0.0001 at two weeks, p = 0.0013 at four weeks, Friedman test). There is a greater log_2_fold-change value for the BCG-SQ arm compared to Mtb+cure (two weeks) and BCG-IV and Mtb+cure (four weeks), indicating that Esx-1 genes are less required in BCG-SQ vaccinated animals compared to other immune animals (Figure 3C). Across genes in the Mce1 system, there is also a significant difference in essentiality by condition (p < 0.0001 at two weeks, p = 0.0012 at four weeks, Friedman test). A similar trend was seen with Mce1 as for Esx-1, with a significantly greater increase in transposon counts in BCG-SQ as compared to Mtb+cure. Thus, despite the similar degree of protection conferred by our immunization strategies, there are quantitative differences among the surviving bacterial transposon mutants, highlighting the potential for TnSeq to uncover unique bacterial genetic requirements under distinct immune conditions.

### Distinct immunization strategies impose differential selective pressures

Given the differences in essentiality of Esx-1 and Mce1 genes across vaccine conditions, we wanted to more comprehensively evaluate how memory responses induced by distinct immunization strategies impact bacterial genetic requirements. In each immune condition, between 4 and 41 genes were conditionally more essential, while between 23 and 61 genes were conditionally less essential (Supplemental Table 1). The less essential genes were highly overlapping across conditions, and the overlap set was largely comprised of genes in the Esx-1 and Mce1 systems (17/23 genes at two weeks, 12/14 genes at four weeks, Supplemental Figure 4A). Six additional genes were less essential in all conditions at the two week time point, including the non-ribosomal peptide synthetase *nrp*/*Rv0101*; *pitA/Rv0545c*, a low-affinity phosphate importer; *Rv0485*, which regulates a PE/PPE pair; the biotin synthesis gene *bioA*/*Rv1568*; *cpsA*/*Rv3484*; and *satS*/*Rv3311*, a protein export chaperone^44^. At four weeks, in addition to the Esx-1 and Mce1 genes, *nrp*/*Rv0101* and *satS*/*Rv3311* remain significantly less essential. Overall, genes that were uniquely less essential in one condition had transposon insertion count log_2_(fold-change) values that trended in the same direction across the other immune conditions, but did not meet criteria for significance, resulting in high correlation coefficients across these genes (Supplemental Figure 4B, C). Together, this suggests a conserved signature in the conditionally non-essential genes under memory immunity, driven by the decreased requirement for the key virulence pathways encoded by Esx-1 and Mce1 genes and other genes known to be essential during acute infection, including *cpsA/Rv3484* and *nrp/Rv0101*^45,46^.

However, as shown in UpSet plots, numerous genes are uniquely more essential under individual immune conditions (Figure 4A). These genes exhibit both quantitative and qualitative differences in transposon insertion count, resulting in lower correlation coefficients when the set of genes more essential in at least one condition are considered (Supplemental Figure 4D, E). For example, *sseA/Rv3283* is significantly more essential in BCG-SQ vaccinated mice four weeks post-infection, non-significantly less essential in BCG-IV and Mtb+cure mice, and unchanged in essentiality in ΔLprG vaccinated mice. *sseA/Rv3283* is part of a membrane-associated oxidoreductase complex, suggesting differences in the oxidative stress experienced by the bacterium in each immune condition^47^. Indeed, no genes were significantly more essential across all conditions at either time point, consistent with a model that each immune condition imposes distinct stresses. Within each immune condition, comparing across the two time points there was little overlap in the more essential genes (Supplemental Figure 4F), consistent with the analysis of genes more essential under BCG-SQ vaccination (Supplemental Figure 1A). This suggests that the immune pressures vary not just by immunization strategy, but also over time, and may reflect differences in the maturation of the memory response under distinct immune conditions.

**Figure 4.**
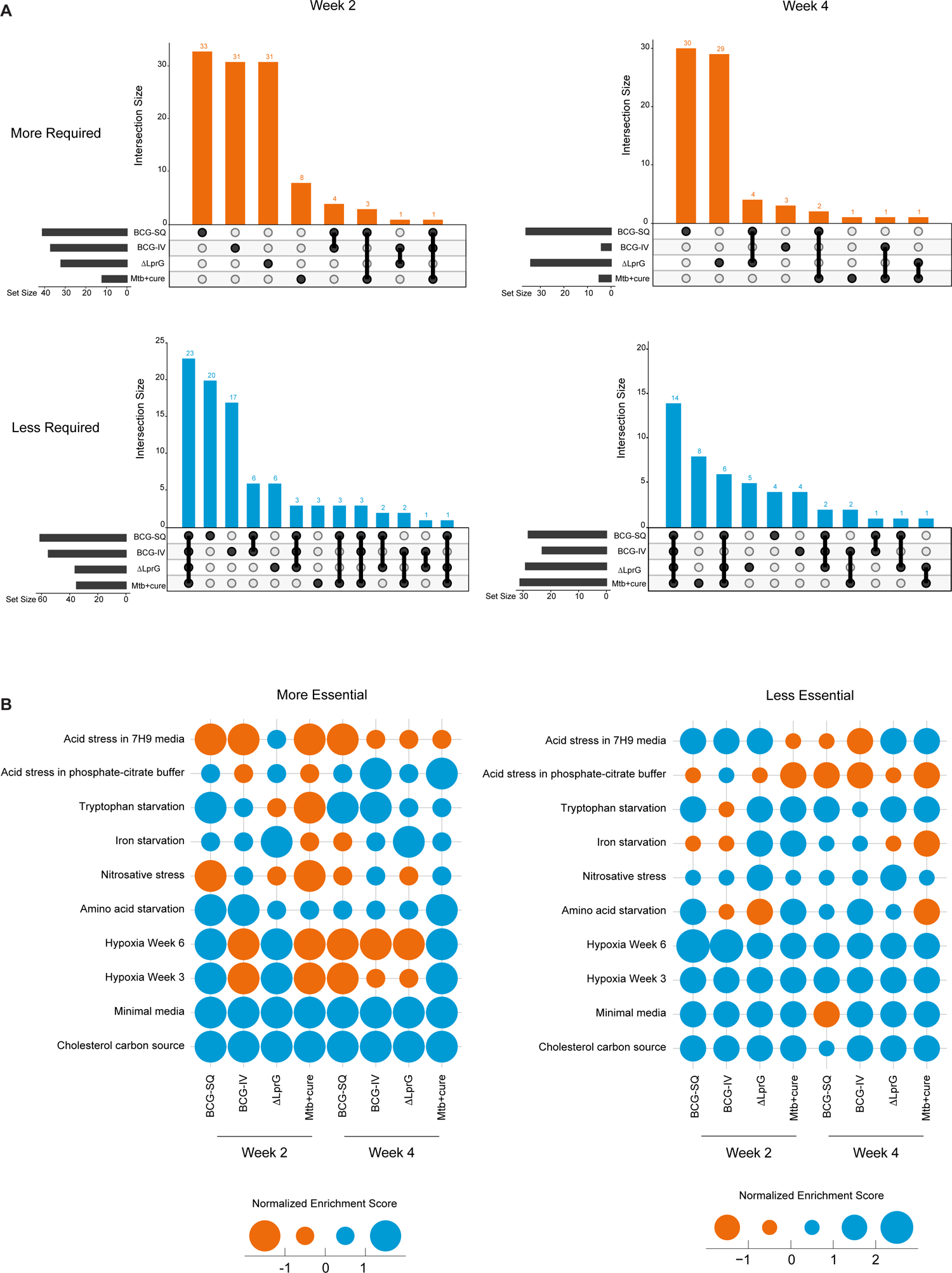
A) UpSet plots of more essential genes in each immunization condition relative to naïve mice (upper panels) and less essential genes in each immunization condition relative to naïve mice (lower panels). B) Bubble heatmaps of GSEA analysis of differentially essential genes. Each bubble represents the enrichment of a gene set, defined by significantly more essential or less essential genes from an *in vitro* condition in the MtbTnDb, against the ranked list of log_2_(fold change) ratio of the TnSeq immunization condition using the GSEA preranked tool. Size of the bubble corresponds to the normalized enrichment score (NES); blue bubbles have a positive NES, and orange bubbles have a negative NES.

To gain more biological insight into the stresses imposed by each immunization strategy and how they change over time, we created gene sets from the defined *in vitro* stress conditions from MtbTnDb presented in Figure 2B. Each set represents genes that are significantly more or less essential under each *in vitro* condition. We then used these sets to perform gene set enrichment analysis (GSEA) against the genes for each immunization arm of our screen, rank-ordered by log_2_(fold-change) in transposon count relative to naïve animals (Figure 4B). This analysis finds that for some *in vitro* conditions, the immune conditions have similar gene essentiality profiles, as indicated by a similar normalized enrichment score (NES). For example, all immune conditions have a positive NES of similar magnitude for the gene sets more essential for growth in cholesterol media and in minimal media, indicating that genes essential for growth in these conditions are not required by the bacterium under immune memory. This suggests that the metabolic milieu experienced by the bacteria is similar across immune conditions.

By contrast, for many of the *in vitro* stress conditions, including acid stress, hypoxia, iron limitation, nitrosative stress, and amino acid starvation, the NES varies both quantitatively and qualitatively (positive *v.* negative enrichment) across immune conditions and over time. For example, genes that are more essential under *in vitro* nitrosative stress are more essential in BCG-SQ vaccinated mice and Mtb+cure mice, as indicated by a large, negative NES at the two week time point. However, the NES for this same gene set is less negative in ΔLprG vaccinated mice and slightly positive in BGC-IV vaccinated animals at this time point. Comparing the two week and four week time points, the NES for this gene set becomes less negative in BCG-SQ vaccinated mice, suggesting a reduction in nitrosative stress over time, and flips from negative to positive in Mtb+cure animals, suggesting an absence of nitrosative stress in this condition. Taken together, this analysis suggests that a TnSeq approach can provide insight into how different immunization strategies achieve bacterial control.

## DISCUSSION

Tuberculosis will remain a global health emergency until a more effective vaccine is available. In the quest to develop an improved TB vaccine, numerous studies have characterized the host response to BCG and new candidates. Despite intensive study, immune correlates of protection remain elusive, and the features that determine bacterial control are incompletely characterized. Here, we have investigated this question from the bacterial perspective, using a functional genomics approach to interrogate bacterial physiology under the selective pressure of adaptive immunity.

Notably, we found that genes encoding the canonical virulence factors Esx-1 and Mce1 are less essential in mice with immunity induced by vaccination or prior Mtb infection. Esx-1 is critical to establishing infection, playing myriad roles including inhibition of phagosome-lysosome fusion; phagolysosomal permeabilization and subsequent stimulation of Type I interferons via the cGAS-STING pathway and activation of the inflammasome via NLRP3; and recruitment of macrophages to form the nascent granuloma, providing a niche for bacterial replication^48–50^. The importance of these, and other, functions of Esx-1 in Mtb pathogenesis is illustrated by BCG, which is avirulent because it lacks Esx-1 due to a large genomic deletion^51^. The Esx-1 secreted antigens ESAT-6 and CFP-10, encoded by *esxA*/*Rv3875* and *esxB*/*Rv3874*, respectively, in addition to being virulence determinants, are immunodominant antigens in mouse and human. We therefore considered whether the relative advantage of Esx-1 transposon mutants in BCG vaccinated mice could reflect a lack of antigenic selection, since these genes are absent in BCG. However, our finding that Esx-1 genes are also less essential in mice vaccinated with the ΔLprG strain, which possesses an intact Esx-1 system, or prior infection with wild-type H37Rv is not consistent with antigenic escape. Rather, our data suggest that one or more of the physiologic roles fulfilled by the Esx-1 system is no longer required in hosts with adaptive immunity.

Many of the roles played by Esx-1 are critical during the initial stages of infection, hinting that vaccination or prior infection recapitulates some aspects of the post-primary infectious milieu, when Esx-1 is no longer needed. Consistent with this interpretation, Mce1 is also important during the early stages of infection. Mce1 was identified as essential during early Mtb infection in the first *in vivo* transposon mutagenesis study by Sassetti *et. al*^53^. In this work, the growth defect of Mce1 transposon mutants resolved later in infection, when Mce4 genes, which encode a cholesterol import system, became essential. This led to the model, since validated, that Mtb consumes fatty acids during acute infection and switches to cholesterol as a nutrient source during chronic infection^54,55^. Our finding that Mce1 is not essential in vaccinated or previously infected mice suggests that some aspects of the chronic infection milieu are recapitulated in immune animals. However, we did not find a concomitant increase in the essentiality of Mce4 genes, suggesting that bacteria surviving the memory immune response utilize another nutrient source, or enter a slowly replicating state. In agreement with our data, a TnSeq study of the genetic requirements for BCG in BCG vaccinated cattle also found that some genes in the Mce1 system became less essential in the context of vaccination, and also did not find that Mce4 genes become more essential^58^. Intriguingly, our data may also suggest a functional link between Mce1 and Esx-1, consistent with comparative genomic analyses^59^. A plausible model for a functional relationship between Esx-1 and Mce1 could be that Esx-1 is required to permeabilize the phagolysosome, permitting access to the host-derived fatty acids imported by Mce1.

Immune correlates of protection following Mtb infection and in the context of vaccination are still lacking^60^. A key finding of this work is that different vaccination strategies induce qualitatively distinct bacterial genetic requirements for survival, and these are different from the bacterial requirements to survive immunity conferred by prior Mtb infection. We interpret the differences in genetic requirements to reflect different immune pressures. Currently, the pipeline for TB vaccine development contains a number of candidate live attenuated vaccines (LAV)^64,65^, with MTBVAC being the most advanced in human clinical trials. Emerging data suggest that there are trade-offs between efficacy and safety for LAV^66^. Our data also support the notion that LAV will differentially mold the memory response and subsequent immunological pressure on the bacterium. Understanding how and why this occurs could be used for rational vaccine design including adjuvant design.

A limitation of this work is that we only investigated one vaccine modality, live attenuated vaccines, and we did not systematically investigate the effect of vaccination route. Future studies to determine whether other vaccine modalities including vectored vaccines and mRNA vaccinations, in additional to vaccine route, also differentially exert immune pressure on Mtb physiology during the adaptive immune response are needed. However, this work highlights the utility of functional genetic screens under vaccine conditions for identifying potential bacterial vulnerabilities that may be targeted in vaccine design. Futures studies using targeted genetic mutants under vaccine conditions may further elucidate how Mtb persists in vaccinated hosts and help explain how and why re-infection occurs.

## METHODS

### Bacterial strains

Unless otherwise noted, Mtb was cultured in Middlebrook 7H9 salts supplemented with 10% oleic acid-albumin-dextrose-catalase (OADC), 0.5% glycerol, and 0.05% Tween 80 at 37°C with agitation, or plated on 7H10 agar supplemented with 10% OADC, 0.5% glycerol, and 0.05% Tween 80 at 37°C. The saturated H37Rv transposon library was previously generated^68^ and outgrown on 7H10 agar supplemented with 10% OADC, 0.5% glycerol, 0.05% Tween 80, 0.2% Cas-amino acids (Difco) and 20 µg/mL kanamycin at 37°C. Bacillus Calmette-Guerin (BCG) was obtained from the Statens Serum Institute and prepared as previously described^69^.

### Animals & infections

Six to eight week old female C57BL/6 mice were purchased from Jackson Laboratories (Bar Harbor, ME.). Infected animals were housed in Biosafety Level 3 (BSL3) facilities under specific-pathogen-free conditions at the Harvard T. H. Chan School of Public Health. The protocols, personnel, and animal use were approved and monitored by the Harvard University Institutional Animal Care and Use Committee.

For BCG and ΔLprG vaccination experiments, animals were vaccinated by injecting 100 µL of frozen bacterial culture (2e7 CFU total) either subcutaneously (BCG and ΔLprG) in the flank or intravenously (BCG) in the lateral tail vein. Mice were rested for 12 weeks post-vaccination prior to infection with the transposon library or single strain infections. For aerosol infections, animals were infected with H37Rv using a Glas-Col instrument (Terra-Haute, Indiana) to deliver approximately 100 CFU. After four weeks of infection, animals were treated with rifampicin (0.2 g/L) and isoniazid (0.1 g/L) in the drinking water for seven weeks. The antibiotic-containing water was changed weekly. Animals were rested for one week to ensure no carry-over effects of antibiotic selection on the challenge transposon library. Transposon library infections were performed by injecting 2e6 CFU of frozen library in the lateral tail vein. Single strain infections with wild-type H37Rv or Esx-1 mutant were performed by injecting 2e6 CFU of mid-log bacterial culture in the lateral tail vein.

### Transposon library sequencing and analysis

At the indicated time points, animals were sacrificed, spleens harvested, homogenized, and plated to recover 1e6 CFU. After three weeks, of outgrowth, colonies were scraped and genomic DNA extracted. Transposon junction sequencing was performed on the recovered genomic DNA essentially as previously described^68^. Read mapping and statistical comparisons of read counts between conditions were performed using Transit v3.2.7. Reads were normalized with the TTR method, insertions in the central 90% of each gene were considered, and a LOESS correction was performed. Repetitive regions, such as PE/PPE genes, were excluded as previously described^68^; excluded genes are listed in Supplemental Table 1.

### Data analysis

Statistical tests were performed with GraphPad Prism v10.3.0. Pathway enrichment analysis was performed with g:Profiler^70^ using Sanger roles for functional annotation. The Sanger terms “IS1081”, “Phage related functions”, “PE_PGRS subfamily”, “PE Subfamily”, “IS6110”, and “PPE family” were excluded from analysis, as were any genes excluded from Transit analysis, as detailed above. Gene set enrichment analysis was performed using the preranked tool^71^. TnSeq datasets were downloaded from MtbTnDb.app on January 25, 2024, and conditions and references are included in Supplemental Table 2. Venn diagrams were generated with Deep Venn^72^. UpSet plots were generated with UpSetR, v1.4.0^73^.

## Data availability

All relevant data to support the findings of this work are located within the paper and supplemental files.

## ACKNOWLEDGEMENTS

We thank Larry Pipkin and Noman Siddiqi, who manage the Harvard T.H. Chan School of Public Health BSL-3 facilities; Shoko Wakabayashi and Jaimie Sixsmith for assistance with mouse experiments; Xin Wang for assistance with microbial cultures; members of the Fortune, Rubin, and Bryson labs for feedback and thoughtful discussions. The support and resources from the Center for High Performance Computing at the University of Utah are gratefully acknowledged. This work was supported by National Institutes of Health grants P01 AI132130 (S.M.F.); T32 AI007061, T32 CA009216, and K08 AI139339 (A.F.C.); K08 AI135098 (A.J.M.); K.S.J. is supported by DP2 AI171122-S1.

## SUPPLEMENTAL INFORMATION

Figure S1. A) Sanger classification of genes that are significantly more essential in BCG vaccinated mice as compared to naïve mice at the two week and four week time points (top panel), and Venn diagrams showing overlap in these gene sets (bottom panel). B) Sanger classification of genes that are significantly less essential in BCG vaccinated mice as compared to naïve mice at the two week and four week time points; p-values annotate significantly enriched Sanger categories as determined by g:Profiler (top panel). Venn diagrams showing overlap in these gene sets (bottom panel). Genes in the overlap set that belong to the Esx-1 or Mce1 pathways are color-coded purple or green, respectively.

Figure S2. A) Schematic of Esx-1 and Mce1 operons illustrating essentiality of each gene at two or four weeks post-infection, relative to naïve animals. B) Plots of transposon junction read count across Esx-1 and Mce1 genes relative to the input library over time in naïve and BCG vaccinated mice. C) Schematic of single strain infection experiment (left) and difference in bacterial burden in naïve as compared to vaccinated animals for wild-type H37Rv and an Esx-1 H37Rv deletion strain in spleen and lung tissues (right). Statistical significance determined by unpaired T-test.

Figure S3. Bacterial burden (CFU) from transposon library challenge experiments.

Figure S4. A) Venn diagrams showing overlap in the less essential genes at two weeks (left panel) and four weeks (right panel) post-infection in immune relative to naïve animals. Genes in the overlap set that belong to the Esx-1 or Mce1 pathways are color-coded purple or green, respectively. B) Heat maps showing log_2_(fold-change) values for all genes that are significantly less essential in at least one of the immune conditions, relative to naïve animals, at two weeks (left) and four weeks (right) post-infection. C) Heat maps showing Pearson correlation coefficients among immune conditions for the less essential genes. D) Heat maps showing log_2_(fold-change) values for all genes that are significantly more essential in at least one of the immune conditions, relative to naïve animals, at two weeks (left) and four weeks (right) post-infection. E) Heat maps showing Pearson correlation coefficients among immune conditions for the more essential genes. F) Venn diagrams showing overlap in the more essential gene sets at two and four weeks for each immune condition.

**Supplemental Table 1.** Transit analysis of transposon junction sequencing data.

**Supplemental Table 2.** List of conditions from MtbTnDb included in Figure 2B and 4B, and citations.

**Supplemental Table 3.** CFU values in the spleen and average log_2_(fold-change) values in transposon read count averaged across the Esx-1 or Mce1 genes across a panel of genetically diverse mice. CFU data were obtained from reference 35, TnSeq data were downloaded from MtbTnDb in order to use standardized data.

