## Supplementary figures and images for "Genetic requirements of *Mycobacterium tuberculosis* for survival under pathogen-specific immunity"

### Supplemental Figure 1

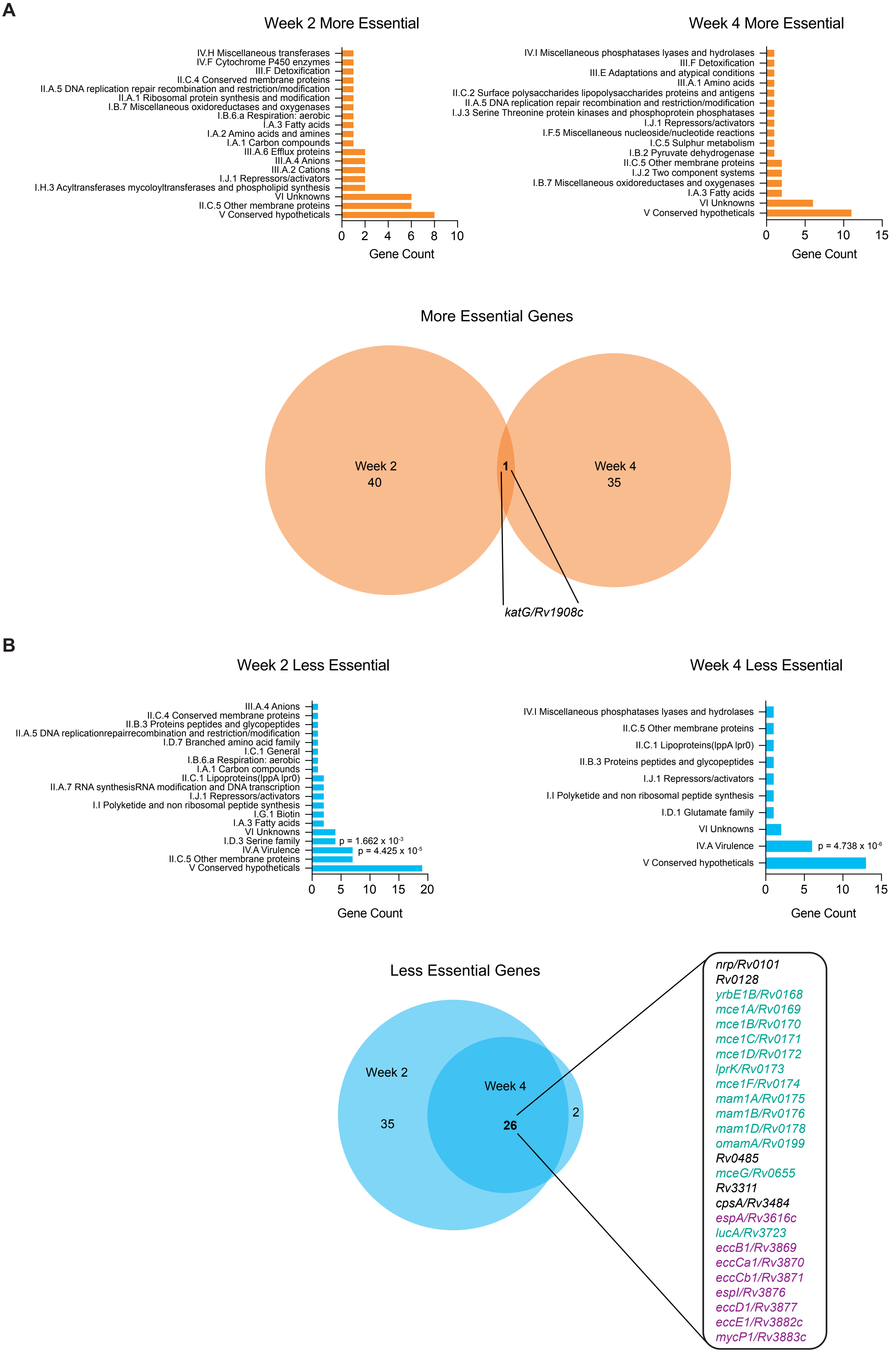

### Supplemental Figure 2

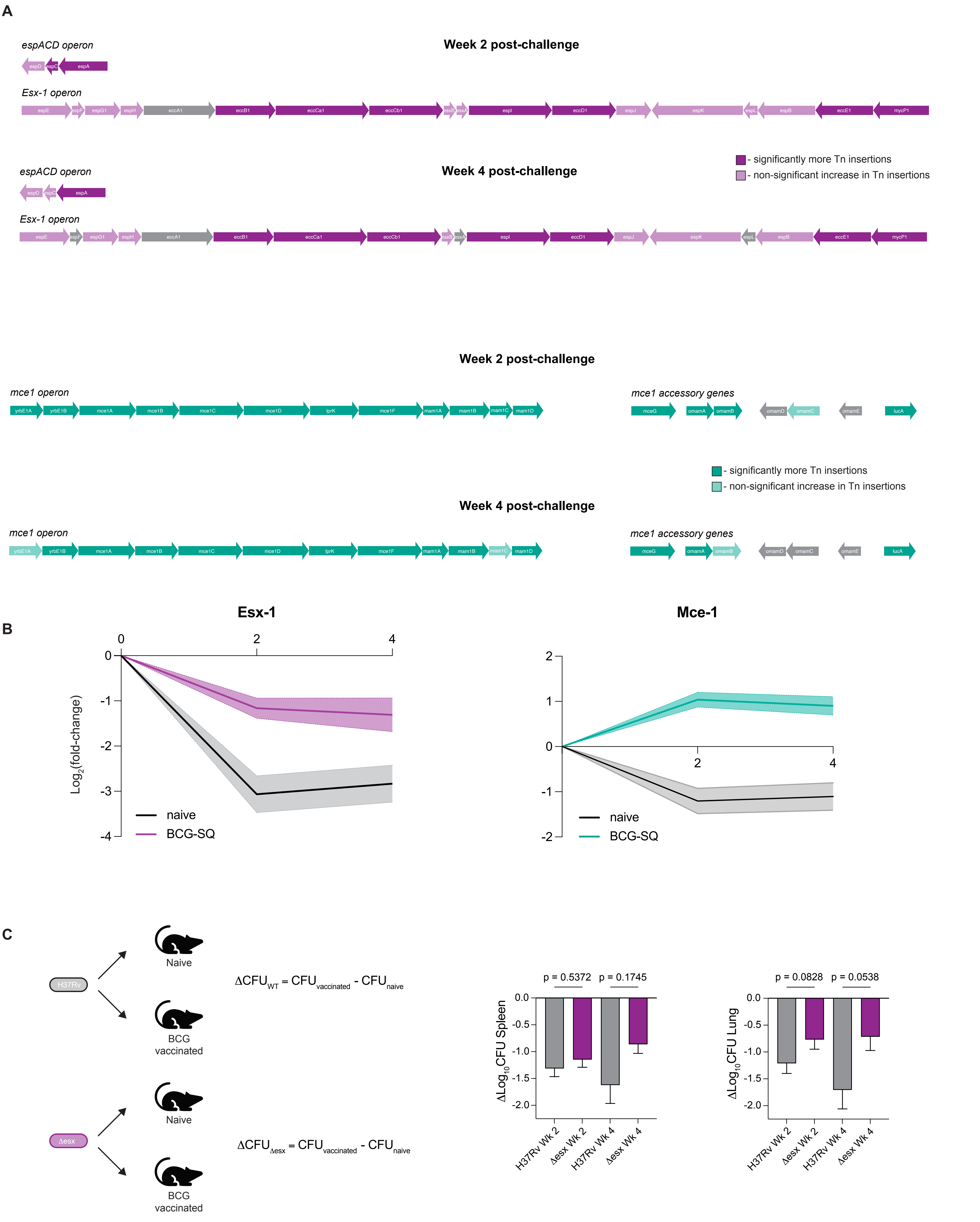

### Supplemental Figure 3

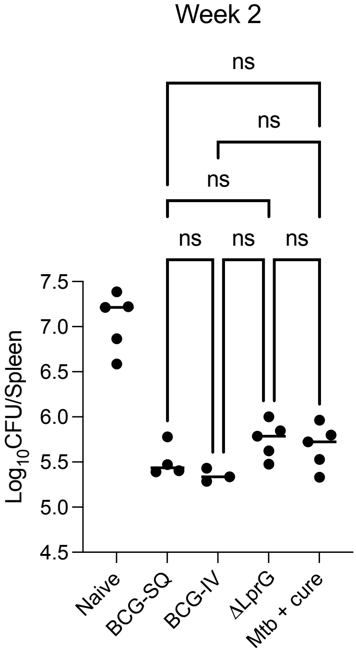

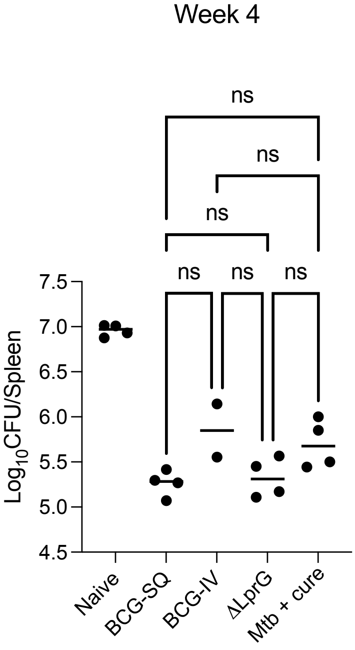

### Supplemental Figure 4

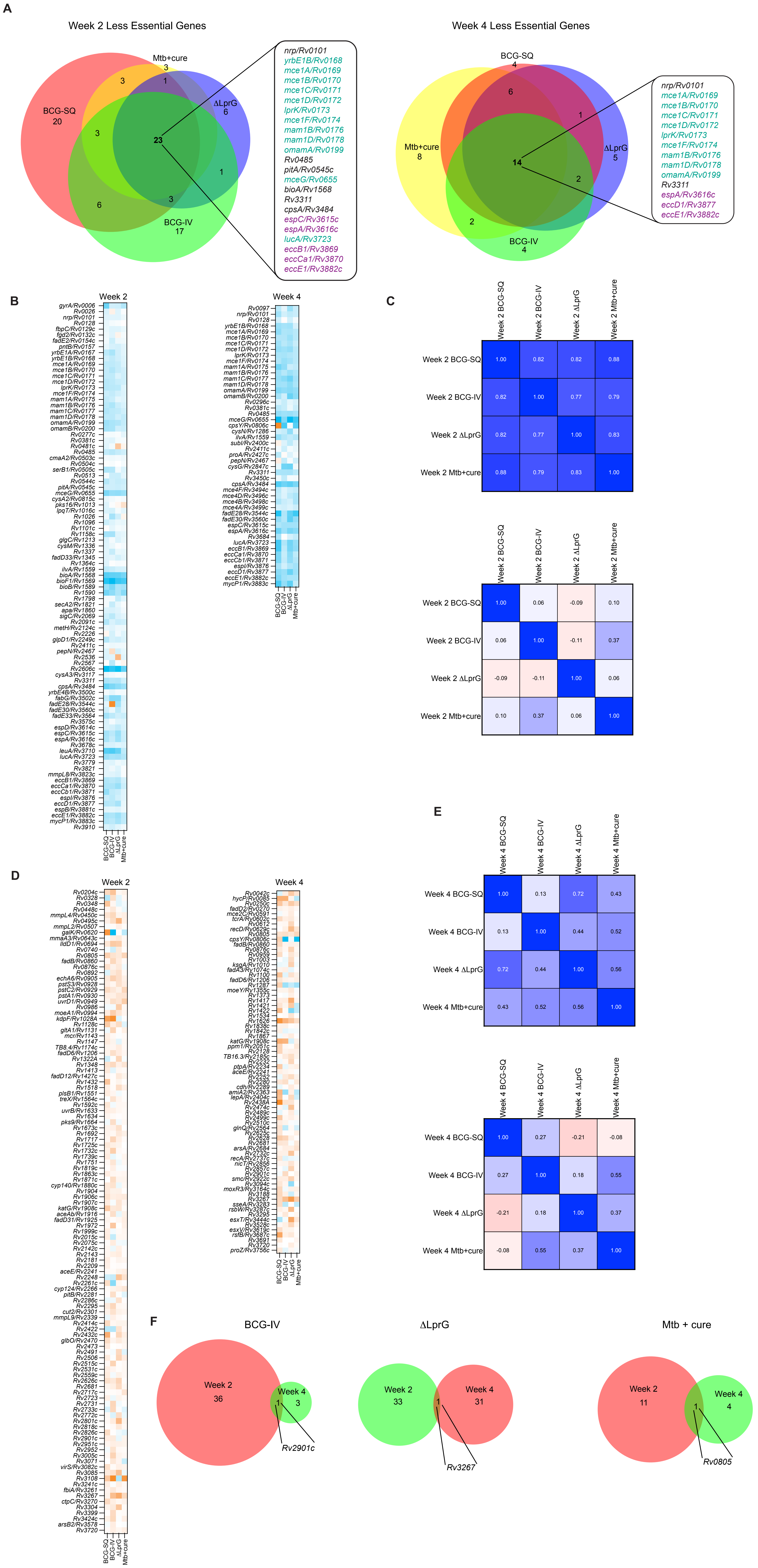
